# Cognitive enrichment improves spatial memory and alters hippocampal synaptic connectivity in a mouse model for early-life stress

**DOI:** 10.1101/2025.08.23.671914

**Authors:** Justin L. Shobe, Elham Ghanbarian, Robert Bain, Rajat Saxena, Meenakshi Chandrasekaran, Bruce L. McNaughton

**Affiliations:** Department of Neurobiology and Behavior, University of California, Irvine, CA, USA; Department of Neurology, University of California, Irvine, CA, USA; Canadian Centre for Behavioural Neuroscience, Department of Neuroscience, University of Lethbridge, Lethbridge, Alberta, Canada; Kavli Institute for Systems Neuroscience and Centre for Algorithms in the Cortex, Norwegian University of Science and Technology (NTNU), Trondheim, Norway

**Author notes:** Please address all correspondence to: Justin L. Shobe, Department of Neurobiology and Behavior, University of California, Irvine, Irvine CA, 92697 USA.

## Abstract

Early life stress (ELS) and enrichment often have opposing effects on long-term cognitive abilities. Deprivation, such as institutionalized care during early childhood neurodevelopmental periods, results in lifelong working memory and recall deficits. In contrast, enrichment facilitates new learning and slows cognitive decline due to aging and neurodegenerative diseases. Similarly, in rodent models, enrichment facilitates learning whereas ELS induces prominent spatial memory deficits. Environmental enrichment (EE) and ELS can cause opposing changes in hippocampal structure (e.g. shifts in synaptic density) that largely depend on experimental conditions. However, it remains untested whether EE can rescue the behavioral disruptions caused by ELS and how this would impact the hippocampus at advanced ages. To address this, we conducted a longitudinal study on ELS mice, extensively training them on a cognitive enrichment track (ET) or an exercise alone control track (CT). After this, the mice underwent repeated memory testing followed by brain extraction for anatomical analysis of their hippocampus. We found that ET reversed spatial memory deficits at 6, 13 and 20 months and reduced the number of dentate gyrus (DG) to CA3 synapses. Surprisingly, this reduction occurred at excitatory MF synapses surrounding CA3 somas in the stratum pyramidale—a layer not typically associated with MF terminals. Collectively, these findings suggest that cognitive enrichment during early adulthood may reverse ELS-induced spatial memory deficits by adjusting synaptic connectivity between the DG and CA3.

## INTRODUCTION

Severe early-life adversity, affecting close to 50% of the world’s children (Fenoglio et al., 2006), can cause both emotional and cognitive disturbances. For instance, children raised in institutionalized settings often exhibit memory deficits and poor impulse control (Pollak et al., 2010); likely contributing to their delayed language acquisition and low scholastic aptitude (Beckett et al., 2007; Eigsti et al., 2011). Evidence suggests that *early* childhood deprivation, such as parental neglect, has a particularly negative impact on executive function and memory (Wodarski et al., 1990; Eckenrode et al., 1993; Spratt et al., 2012) because this sensitive time period (first 2-3 years in humans and first 3-4 weeks in rodents) is critical for the maturation of brain systems necessary for learning and memory, such as the hippocampus (Malave et al., 2022; Kloc et al., 2020; Leinekugel et al., 2002). This may be why ELS during this time period can cause long-lasting deficits in declarative learning and hippocampal function. Adults who have experienced ELS have difficulty remembering episodic events and perform poorly on delayed word recall tasks (Ding and He, 2021; Ma et al., 2021; Cai et al., 2024). These individuals have smaller hippocampal volumes, especially their dentate gyrus (DG) (Humphreys et al., 2019; Youssef et al., 2019; Koyama et al., 2022; Kawamoto et al., 2023). Importantly, these findings have been replicated in rodent models designed to mimic maternal neglect (Walker et al., 2017). As adults, these ELS animals perform poorly on hippocampus-dependent spatial memory tasks such as the Morris water maze and object location memory (OLM) (Brunson et al., 2005; Cui et al., 2006; Rice et al., 2008; Ivy et al., 2010; Pollak et al., 2010; Naninck et al., 2015; Chen et al., 2016; Molet et al., 2016b; Naninck et al., 2017) consistent with human findings, these animals have smaller hippocampi and dendritic atrophy in CA3 and CA1 neurons (Brunson et al., 2005; Molet et al., 2016a; Teicher et al., 2016).

Unfortunately, there are very few non-invasive treatments for individuals suffering from the effects of ELS; however, experience-based behavioral interventions may help. For example, the Bucharest intervention project (and related studies) found that adoption of institutionalized children into more nurturing environments correlated with improved cognition and higher IQ scores (i.e., the earlier the better) (Tizard and Rees, 1974; Nelson et al., 2007; Bos et al., 2009; Almas et al., 2016). Enriching activities later in life can also facilitate cognitive recovery (Cai et al., 2024).

In rodent models, environmental enrichment (EE) improves cognitive functions, including spatial learning, memory, and task learning (Cheng et al., 2022; Rountree-Harrison et al., 2018; Bennett et al., 2006; Williams et al., 2001; Woodcock et al., 2000; Zeleznikow-Johnston et al., 2017). Consistent with this, we found that extensive cognitive enrichment leads to dramatic improvements on a wide variety of memory tasks (Gattas et al., 2022). Mechanistically, EE promotes synaptogenesis and adult hippocampal neurogenesis (Speisman et al., 2013) increases spine count (Jung et al., 2014) and dendritic complexity (Connor et al., 1982), modulates synaptic signaling (Pintori et al., 2024) and enhances long-term potentiation in the hippocampus (Cortese et al., 2018; Artola et al., 2006). Additionally, it improves sensory processing and neural coding efficiency (Engineer et al., 2004; LeMessurier et al., 2019). For instance, EE improves the ability of the hippocampus to distinguish between different environments (i.e., better pattern separation), which in turn promotes better spatial learning (Bilkey et al., 2017; Ventura et al., 2024).

The DG-CA3 connection is particularly susceptible to experience-dependent plasticity (Henze et al., 2000; Urban et al., 2001). ELS and EE can have opposing effects on this such as the rate of neurogenesis in the DG and the magnitude of DG-CA3 long-term plasticity (LTP and LTD) (Alwis & Rajan, 2014; Kempermann, 2019). In contrast, both ELS and EE increase mossy fiber sprouting (Brunson et al., 2001; Galimberti et al., 2006; Bramati et al., 2023). This is especially interesting considering the positive correlation between mossy fiber expansion and spatial memory performance (Schwegler et al., 1990; Ramı rez-Amaya et al., 2001; Carasatorre et al., 2015). With respect to EE, however, it is important to recognize that many of these structural and physiological changes largely depend on experimental conditions such as the animal’s age and the duration of enrichment (Gogolla et al., 2009; Bednarek and Caroni, 2011); making it difficult to predict how ELA and EE would mechanistically interact.

Given these observations, we set out to test the hypothesis that cognitive enrichment would mitigate the negative impact of ELS on learning and determine whether this would be accompanied by structural changes in the mossy fiber terminals, which are primarily considered glutamatergic (Walker et al., 2001; Gutierrez et al., 2003); however, there is some evidence that these terminals also contain GABA (Cabezas et al., 2012; Uchigashima et al., 2007). Using a longitudinal approach, we first trained ELS mice on our complex obstacle enrichment track (ET). This 3-month protocol produces dramatic and broad long-term memory enhancements well above that of standard enrichment procedures (Gattas et al., 2022). Moreover, the incorporation of a simple ramp control track (CT) group allowed us to disambiguate the effect of exercise alone. Following this, we repeatedly tested mice on spatial and object recognition tasks across their lifespan. Finally, we assessed whether these transgenic mice (GCaMP6fs) had hippocampal structural changes using a combination of endogenous fluorescence and nanobodies. To specifically visualize MF projections and their synapses, we took advantage of the selective expression of the GCaMP6fs signal in the DG (no signal in CA3 neurons) together with excitatory synaptic markers such as PSD-95. Unexpectedly, we found clusters of these atypical excitatory MF synapses surrounding CA3 cell bodies whose density was regulated by the cognitive component of enrichment.

## RESULTS

### Cognitive enrichment rescues the spatial memory deficits of ELS mice

First, we induced ELS using the disrupted maternal care paradigm (limited bedding and nesting method, LBN) since it causes progressive memory deficits as rodents age (Molet et al., 2016; Rocha et al., 2021). Following this we assigned mice to either enrichment or control groups (Fig 1A). Our previous findings suggest that ET training produces broad and dramatic memory enhancements (Gattas et al., 2022). We speculated, however, that a scaled back version of the ET training protocol would more selectively benefit hippocampal circuits that are especially vulnerable to ELS (Scholz et al., 2015; Molet et al., 2016b; Hoeijmakers et al., 2018; Manno et al., 2022). Thus, we ran our enrichment protocol for three thirty-minute sessions per week, one quarter of our original six one-hour sessions per week.

**Fig. 1:**
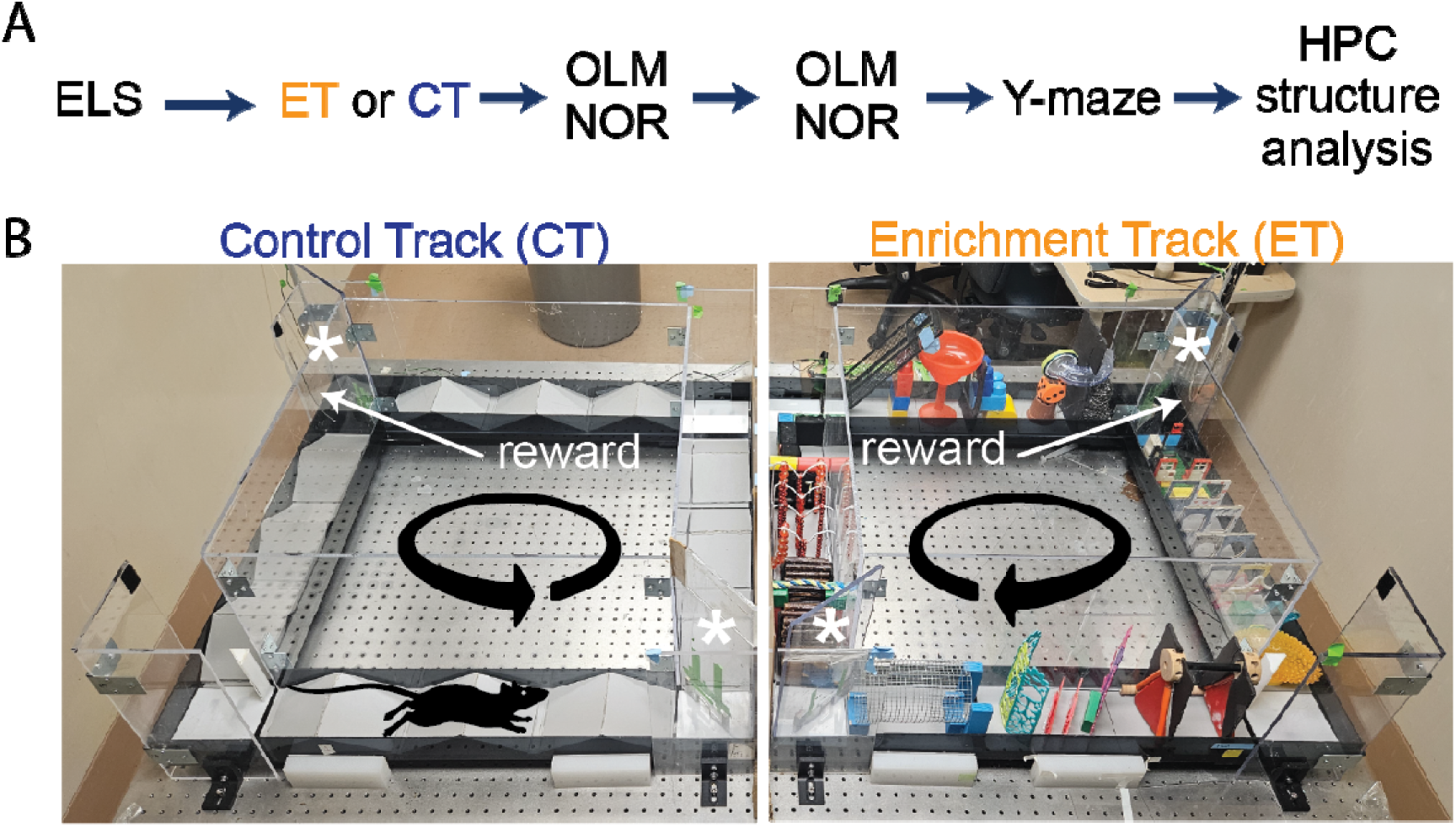
Experimental timeline of enrichment training, behavioral tests and structural analysis on early-life stressed (ELS) and control mice. **A)** Initially, we separated ELS mice into two sex matched groups. Half of the ELS mice (4 males and 4 females) ran on the ET track (orange) while the other half (4 males and 4 females) ran on the CT track (dark blue). Following this, both groups received longitudinal behavioral testing and hippocampal structural analysis. **B)** We ran double-housed adult ELS mice on our automated side-by-side environmental enrichment (EE) setups such that one mouse ran on the control track (CT, left panel) while its cage-mate ran on the enrichment track (ET, right panel). On the ET track, mice had to navigate through multiple obstacles (obstacle configuration changed every session), whereas the CT only ran over simple ramps. We place two one-way doors (asterisks) at opposite corners of the square track to minimize backtracking. Mice received a reward (sweetened condensed milk) upon completion of each lap controlled by an automated delivery system using an overhead camera tracking system.

We then tested both groups (ELS+ET, 4M and 4F, n=8) and (ELS+CT, 4M and 4F, n=8) on OLM and NOR memory tests at mature (6 mo), middle (13 mo) and old (20 mo) ages to determine if behavioral gains would last as the mice aged (Fig. 1A &B). We observed a significant group level (ET vs CT) difference in OLM (three-way ANOVA, F(1,12) = 8.80, p = 0.012) but no significant effect of age (F(1,12) = 1.69, p = 0.218) or sex (F(1,12) = 0.97, p = 0.345) (Fig. 2A). In contrast to OLM and also the results of Gattas et al., 2022, we found no significant group (three-way ANOVA, F(1,12) = 0.074, p = 0.791), age (F(1,12) = 2.93, p = 0.113), or sex (F(1,12) = 0.11, p = 0.744) effect in the NOR test (Fig. 2B). In the final OLM/NOR time-point (18 months), the mice, unfortunately, did not spend enough time exploring the objects to get accurate testing scores (values are the sum of investigation times for both objects). Compared to their mature and middle-age scores, these older mice spent significantly less time investigating the objects (Sup. Fig. 2B, two-way ANOVA, P<0.0001, F(2,88) = 30.86); likely due to habituation to the OLM/NOR setup, an age-dependent reduction in exploratory behavior, or a combination of both. To reinvigorate their exploratory behavior, we switched to a completely novel setup, the spatial Y-maze. On test day, we found that ET mice (2M &3F, n=5) spent significantly more time exploring the novel arm than CT mice (3M &4F, n=7) in a group comparison (two-way ANOVA, F(1,8) = 19.89, p = 0.002), with no significant effect of sex (F(1,8) = 1.81, p = 0.215), suggesting that ET produced a life-long rescue of spatial memory formation or retention (Fig 2C and D)

**Fig 2.**
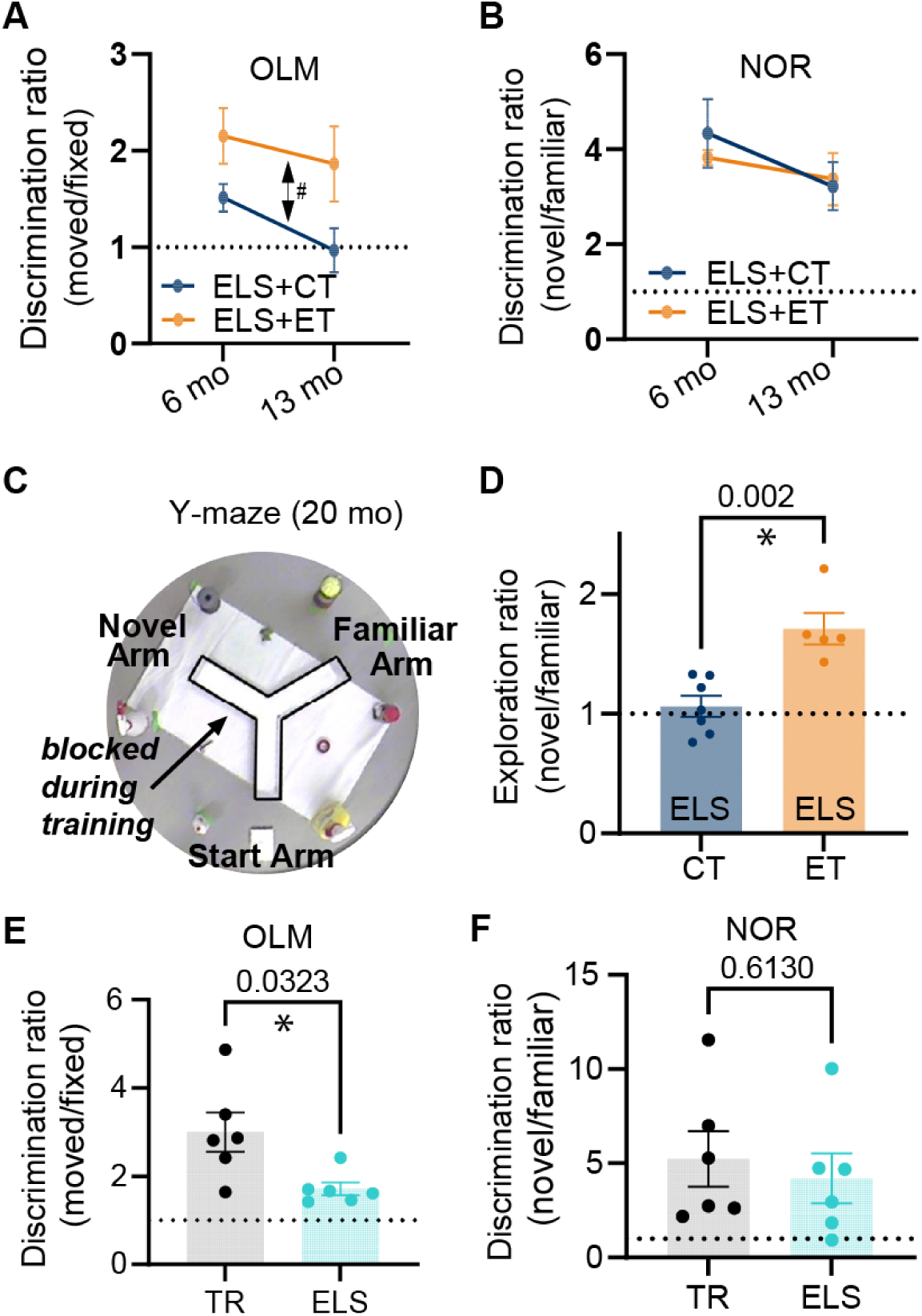
Enrichment track (ET) training reverses life-long spatial memory deficits caused by early-life stress (ELS). Longitudinal testing of ELS mice on object-location (OLM) and novel object recognition (NOR) at mature (6 mo) and middle-age (13 mo) time points revealed that **A**) the ELS+ET group (orange, n=8 mice) had improved 24hr memory (double-sided arrow, three-way repeated measures ANOVA, F(1,12)=8.80, p=0.012) for OLM relative to the ELS+CT control group (dark blue, n=8). There was no significant effect of age or sex. **B**) In contrast for NOR, we found no group difference (F(1,12)=0.074, p=0.791) and no effect of age or sex. The same mice (n=16) were tested on OLM and NOR. A value of 1 represents equal time with both objects (dashed line). **C**) In a final (20 mo) Y-maze test, the surviving mice (5M & 7F, n=12) explored the two open arms for 5 min while the third arm remained blocked (transparent walls surrounded by distinct external cues) followed 24hr later by testing (5 min session) where mice were allowed to freely explore all three arms. **D**) The ELS+ET group (orange, n=5) spent significantly more time exploring the novel arm in comparison to the ELS+CT group (dark blue, n=7) (two-way ANOVA, F(1,8)=19.89, p=0.002). We calculated the active exploration ratios by dividing the occupancy times of the novel arm by that of the familiar arm (we excluded periods of immobility lasting longer than 1 sec). E&F) To verify that ELS alone (in the absence of EE) induced memory deficits, we tested mice on OLM and NOR using the same conditions as group 1. **E**) In OLM, the TR group (black dots, n=6 males) had significantly higher discrimination ratios (moved/fixed) compared to the ELS group (cyan dots, n=6 males), indicating that they spent more relative time investigating the moved object (p=0.0323, unpaired two-tailed t test with Welch’s correction). **F**) For NOR we observed that no significant difference in discrimination ratios between the same TR and ELS groups (p=0.613, unpaired two-tailed t test). Plots include values from individual mice (circles in D-F), mean (bar height and circles in A&B), and SEM (error bars).

Importantly, our design mitigates the exercise confound since both groups must run laps to receive rewards. In fact, the control track (CT) group, that ran over simple ramps, completed more laps than the ET group that had to navigate through complex obstacles (Sup Fig. 1C, unpaired two-tailed t-test, p<0.0001). Thus, additional exercise cannot account for these observed memory enhancements in the ET group.

### ELS causes spatial memory deficits

Many studies have shown that ELS causes significant memory disruptions (Brunson et al., 2005; Cui et al., 2006; Rice et al., 2008; Ivy et al., 2010; Wang et al., 2011; Naninck et al., 2015, 2017; Molet et al., 2016a; Hoeijmakers et al., 2018). To confirm these findings, we tested a separate non-enriched group of male mice under either ELS or typical rearing (TR) conditions (Fig1a, chartreuse arrow). These ELS mice and their age-matched TR controls remained under normal housing conditions until behavioral testing at mature adult ages (∼6 months). The ELS group (6 mo) showed memory impairments in the OLM task (p=0.032, two-tailed t-test, Fig. 2E bottom) but not in NOR (p=0.613, Fig. 2F bottom). Taken together, our findings are consistent with other studies that ELS can cause a selective disruption of hippocampal-dependent memories (Molet et al., 2016b; Hoeijmakers et al., 2018) but also see (Ivy et al., 2010; Naninck et al., 2015).

### Cognitive enrichment does not change gross hippocampal morphology

Studies have shown that enrichment and ELS can cause volumetric changes in hippocampal subfields such as the mossy fiber (MF) pathway (Brunson et al., 2005; Galimberti et al., 2006; Molet et al., 2016b; Teicher et al., 2016; Humphreys et al., 2019; Youssef et al., 2019; Bramati et al., 2023). This prompted us to examine whether these ELS GCaMP-expressing mice mice had any gross morphological changes in their hippocampal structure. We used Thy1-because our earlier pilot images revealed that their DG neurons and axons (the MF pathway) were filled with GCaMP signal. One month following the final Y-maze test (21 months of age), we extracted brains for fluorescent imaging of endogenous GCaMP and DAPI signals in the dorsal hippocampus. Importantly, this GCaMP signal was absent from CA3 neurons, indicating that the GCaMP signal in the CA3 subfield was specific to MFs (Fig. 3B lower right inset). As expected, there was sparse GCaMP expression in CA1 neurons (Fig. 3B top inset).

**Fig. 3:**
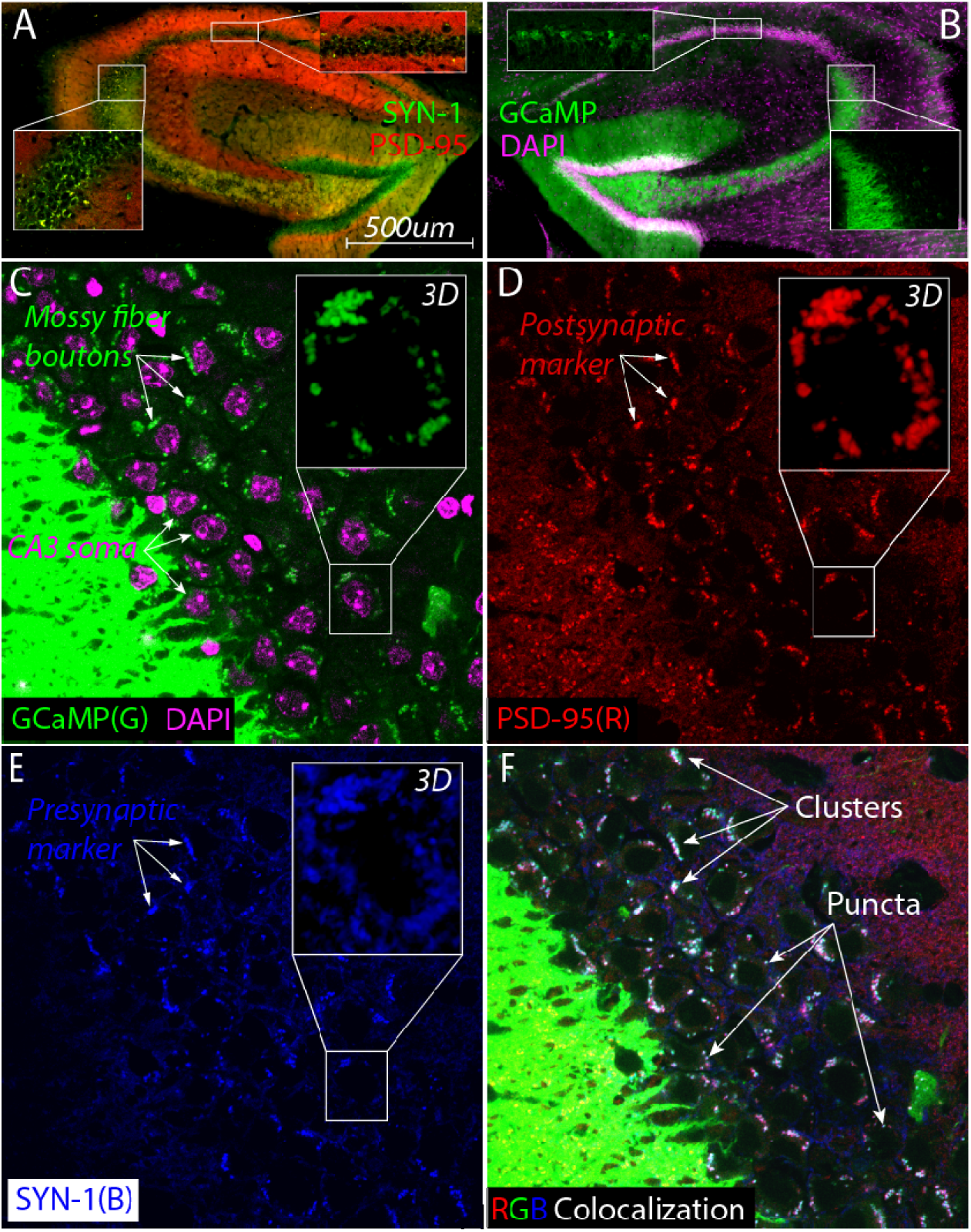
Large synaptic clusters containing mossy fiber boutons surround CA3 cell bodies in aged ELS mice (20 months). 3A) Colocalization of postsynaptic marker DPSD-95 nanobody (red) with presynaptic marker DSynaptotagmin nanobody (SYN1, green) reveals the presence of large synaptic clusters (yellow) around CA3 soma (bottom inset) but not in the CA1 pyramidal subfield (top inset). **3B)** High expression of GCaMP signal (green) in the dentate gyrus cells show that mossy fibers project and appear to terminate near CA3 cell bodies (purple, DAPI). GCaMP signal is not detectable inside CA3 neurons (right inset) but it does fill a large percentage CA1 pyramidal neurons (top inset) as expected from imaging studies. **3C-F)** Super resolution single-plane optical section imaging of the stratum pyramidale (**3C**, CA3 cell bodies labeled with DAPI, purple) stained for mossy fiber boutons (**3C**, green), postsynaptic PSD-95 (**3D**, red) and presynaptic SYN-1 (**3E**, blue) yields extensive triple colocalization (**3F**, white) that consists of larger clusters and smaller individual puncta consistent with the hallmarks of MF synapses. Max Z projections of zoomed in z-stack images from a single cell body (**3C-E,** insets) demonstrate that these synaptic clusters surround the cell in all three dimensions.

Given the specificity of our GCaMP signal in combination with DAPI staining, we measured the areas of the GC Layer, the Hilus, and the suprapyramidal blade of the MF. After normalization to total hippocampal area, we did not detect any significant differences in these areas between ELS+ET (n=5) and ELS+CT (n=7) mice (Sup Fig. 4B). The number of CA3 neurons per 0.01mm² was not significantly different between groups. Altogether, this suggests that ET training does not cause gross structural changes in the DG to CA3 circuit when compared to our exercise alone control group (CT).

**Fig. 4:**
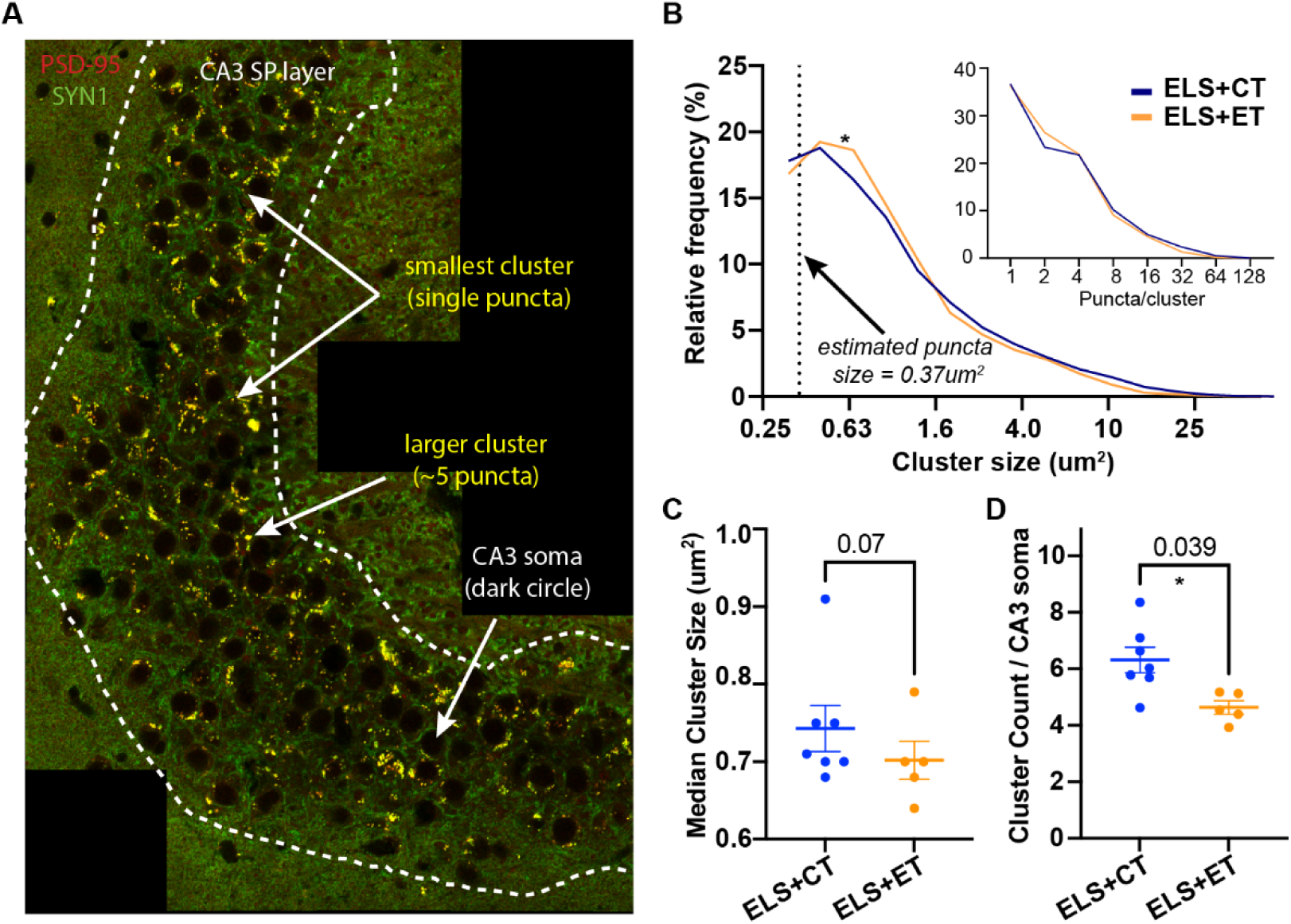
ET training reduces the density of putative MF synapses in the CA3. 4A) A representative tiled image from an ELS+ET mouse (brain harvested following Y-maze) double-stained with PSD95 (red) and SYN1 (green) nanobodies. In the stratum pyramidale (SP) region (dashed white line) of the CA3, colocalized (yellow) clusters appear to be comprised of individual puncta (area = ∼200 pixels or 0.37um^2^). Clusters smaller than 150 pixels were removed from subsequent analysis. **4B**) Histogram of log-transformed cluster sizes (pooled) and estimated number of puncta/cluster (inset) identified from ELS+ET (10,497 clusters, n=7 mice) and ELS+CT mice (5847 clusters, n=5 mice). Statistical analysis between the groups using rank-based (Mann Whitney, p=0.035) and cumulative (Kolmogorov-Smirnov, p=0.0001) methods indicate significant differences (asterisk) in their distributions, likely due to fewer puncta per cluster in ELS+ET mice (inset). Dashed line indicates the x-intercept corresponding to our estimated size of a single puncta. **4C&D**) A mouse-by-mouse analysis reveals that ELS+ET mice (n=5, orange dots) have fewer clusters compared to ELS+CT controls (n=7, dark blue dots, two-way ANOVA, F(1,8)=6.07, p=0.039, asterisk, **4D**). There was a trend for smaller median cluster sizes in the ELS+ET group (two-way ANOVA, F(1,8)=4.36, p=0.070, **4C**). Tiled sections were normalized to counts to the total number of CA3 neurons.

### Identification of atypical MF synapses surrounding CA3 cell bodies

To visualize synapses, we found it necessary to use nanobodies that are capable of penetrating fixed tissue (Kilisch et al., 2023) allowing us to compare presynaptic and postsynaptic markers as well as the endogenous GCaMP signal (the MF indicator) in the same slice. Immunostaining with synaptotagmin and PSD-95 revealed the presence of prominent large synaptic clusters surrounding the CA3 soma that were not as visible in other areas such as the CA1 subfield (Fig. 3A top inset).

The large synaptic clusters, consistent with MF boutons, were particularly notable in the CA3 stratum pyramidale, which was unexpected. To confirm these were bona fide MF synapses, we individually visualized all three signals: mossy fiber GCaMP (Fig. 3C, green), postsynaptic PSD-95 (Fig. 3D, red), and presynaptic synaptotagmin (Fig. 3E, blue). Around CA3 cells (purple staining in Fig. 3C), we observed many clusters in the same location across all three images (white arrows in Fig. 3C-3E). The observed putative MF boutons colocalized with PSD-95 (Fig. 3C and 3D) as well as VGLUT1 (Sup Fig. 5) suggesting that they were excitatory synapses.

**Fig. 5:**
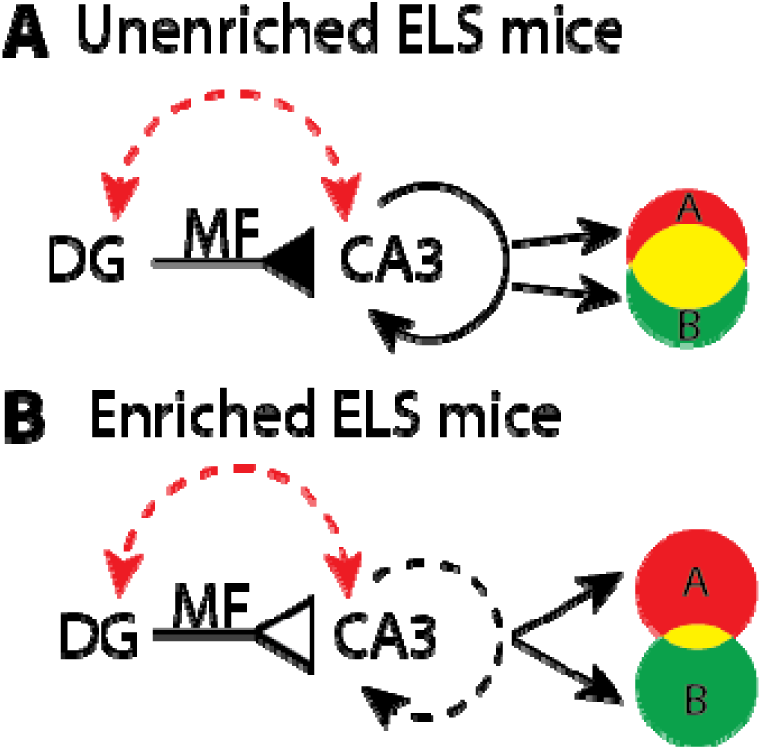
Hypothetical model that illustrates how enrichment could improve pattern separation in aged ELS mice. A) Age and stress (red dashed arrow) increase the excitability of DG and CA3 which may cause mossy fiber (MF) input (solid inverted arrow) to drive dense recurrent activity (solid circular arrow). These dynamics could produce large overlaps (yellow shading) between context specific representations (A and B). **B)** In enriched mice, age and stress may still lead to high levels of DG and CA3 excitability. However, a 27 percent reduction in the number of atypical MF synapses, per CA3 neuron, (open inverted arrow) could help to restore pattern separation by driving sparser recurrent activity (dashed circular arrow) that orthogonalizes (less yellow shading) context specific representations (A and B).

The triple colocalization image (overlay of all three RGB colors) revealed that many clusters were white, suggesting they were composed of all three markers (Fig. 3F). These clusters varied in size, from larger clusters (top three arrows, Fig. 3F) to individual puncta (bottom three arrows, Fig. 3F). We performed a 2D spatial cross-correlation on these images to statistically determine the extent of colocalization. A pairwise analysis of all three combinations (PSD95-SYN1, PSD95-GCaMP, and SYN1-GCaMP) revealed significant overlap, with peaks higher than the shuffled distribution (Sup Fig. 3). This statistically significant triple colocalization of mossy fiber boutons and pre- and postsynaptic markers strongly supports the presence of excitatory MF synapses in the CA3 pyramidal layer.

### Characterization of atypical MF synapses in the SP layer

Next, we examined quantitative differences in putative excitatory MF synapses between the ELS+ET and ELS+CT mice. We immunostained sections for postsynaptic PSD-95 (red) and presynaptic synaptotagmin 1 (SYN1, green) as this combination of synaptic markers had the highest overlap score (see the green line, Sup Fig. 3). We tiled a large area of the CA3 bend to identify MF synapses (yellow clusters and white arrows) of various sizes as well as CA3 cell bodies (dark circles) (Fig. 4A). Regardless of size, colocalized clusters in the stratum pyramidale (SP) region (dashed white line) of the CA3, appeared to be composed of individual puncta with an area = ∼200 pixels or 0.37 µm² (Fig.4 A).

We used automated counting to determine frequency and size of clusters in the tiled images (2 per mice) between ELS+ET and ELS+CT mice. In total, ELS+ET mice had 10,497 clusters (n=7 mice) while ELS+CT mice had 5,847 clusters (n=5 mice). Statistical analysis between the pooled groups using rank-based (Mann-Whitney, p=0.035) and cumulative (Kolmogorov-Smirnov, p=0.0001) methods indicated significant differences in their size distributions. ELS+ET mice (orange line) had fewer large clusters and proportionally more smaller clusters than ELS+CT controls (blue line, Fig. 4B). ELS+ET mice had a corresponding shift in puncta per cluster in (Fig. 4B inset); assuming that the larger clusters are composed of discrete puncta.

In a mouse-by-mouse analysis, normalized per CA3 neuron, we found that ELS+ET mice had on average 27% fewer clusters (4.75±0.15, n=5, orange dots) compared to ELS+CT controls (6.31±0.63, n=7, dark blue dots, two-way ANOVA, F(1,8)=6.07, p=0.039) with no significant sex differences (F(1,8) = 0.043, p = 0.841 Fig. 4D). There was no significant group or sex difference in median cluster size (two-way ANOVA, group: F(1,8)=4.36, p=0.070; sex: F(1,8) = 0.0009, p = 0.976, Fig. 4C), although there was a small trend for smaller synapses in ELS+ET mice consistent with our distribution analysis. These findings suggest that ET training reduces the number of mossy fiber (MF) synapses surrounding CA3 somas in ELS mice without significantly affecting the size of individual synaptic clusters.

## DISCUSSION

In a mouse model of ELS, we found that cognitive enrichment training during young adulthood leads to long-lasting changes in spatial memory and hippocampal structure. To our knowledge this is the first demonstration of these effects in a rodent model of maternal neglect. Compared to exercise alone controls (ELS+CT), mice that underwent enrichment training (ELS+ET) showed significant improvements in long-term OLM (24hr) memory at both mature (6 months) and middle-ages (13 months). Additionally, at 20 months, ELS+ET mice performed better memory in a final spatial Y-maze test. However, we found no group differences in NOR, a task which relies less on the hippocampus (Oliveira et al., 2010; Barker and Warburton, 2011; Cohen and Jr., 2015). Surprisingly, our anatomical analysis of the hippocampus revealed that aged ELS mice had prominent MF-associated excitatory clusters surrounding the CA3 neurons, an area not typically associated with MF synapses. Furthermore, we found that in aged ELS mice early enrichment reduced the number of these atypical MF synapses by ∼25%.

An important caveat in our findings is that within the stratum pyramidale, we cannot distinguish between atypical excitatory mossy fiber boutons that synapse onto pyramidal cells versus those that synapse onto inhibitory interneurons (Henze et al., 2000; Urban et al., 2001). However, excitatory pyramidal neurons vastly outnumber inhibitory interneurons in the CA3 pyramidal layer, with interneurons comprising only approximately 10-15% of the total neuronal population (Freund and Buzsáki, 1996; Klausberger and Somogyi, 2008). This numerical predominance suggests that the reduction in excitatory mossy fiber synaptic clusters we quantified likely represents connections onto pyramidal cells. Nevertheless, given that interneurons in this layer provide feed-forward inhibition that regulates CA3 excitability (Jinde et al., 2012), and that environmental enrichment has been shown to modulate inhibitory neurotransmission (Speisman et al., 2013; Cortese et al., 2018), changes in mossy fiber connectivity onto inhibitory neurons could alter CA3 pattern separation. Future studies using cell-type-specific markers or electrophysiological approaches will be necessary to determine whether cognitive enrichment and/or ELS differentially alters these synapses.

There are important similarities and differences between our behavioral observations and those of previous studies. Earlier studies found that ELS can cause wide-ranging spatial (e.g. OLM) and recognition (e.g. NOR) memory impairments (Brunson et al, 2005; Hoeijmakers et al., 2018; Ivy et al., 2010; Molet et al., 2016; Naninck et al., 2017; Bolton et al., 2020; Short et al., 2020; Rice et al., 2008). These impairments; however, appear to occur at different rates during aging. A careful study, in rats, found that spatial memory deficits start during adolescence whereas the effects of ELS on NOR memory does not occur until older ages (Molet et al., 2016) (however see male mice in Naninck et al., 2015 study). Similarly, we found that our ELS alone mice (6 mo) had normal NOR memory but significant deficits in OLM, consistent with the model that the hippocampus is more susceptible to assaults from ELS than other brain regions. Interestingly our observations also suggest the ET protocol has a more selective effect on hippocampus function because between ET and CT mice we found no group differences in NOR. This finding seemingly contrasts with our initial use of the ET protocol, where we found broad memory enhancements (Gattas et al., 2022). The original study; however, used a more extensive protocol (six 1-hour sessions/week versus three 30-minute sessions/week) that started at an earlier age. This suggests that the abbreviated ET protocol used in this study produces a more limited effect that is biased towards improvements in hippocampal function.

Contrary to our predictions, we found no differences in MF volume between the ELS+ET and ELS+CT groups. Other studies, conducted with non-stressed mice, report that enrichment causes mossy fiber expansion beyond the SL layer (Galimberti et al., 2006; Bramati et al., 2023). Unlike our design, those studies did not control for exercise, as only the EE group had access to a running wheel. Since exercise alone contributes to mossy fiber sprouting (Toscano-Silva et al., 2010), it may be exercise, rather than cognitive enrichment, that drives the growth of mossy fibers toward the CA3 cell bodies.

Several factors could explain why these synapses remain uncharacterized, despite their previous observation (Amaral and Dent, 1981; Dailey et al., 1994; Qin et al., 2001; Banks et al., 2024). Age and ELS cause CA3 dendritic atrophy and mossy fiber sprouting (Brunson et al., 2001; Galimberti et al., 2006; Adams et al., 2010; Molet et al., 2016b), which may be an attempt by the MFs to compensate for synaptic loss, shifting synapse density toward CA3 cell bodies. Another possibility is our use of nanobodies, which are ∼10 times smaller than antibodies. This increases their ability to detect antigens in fixed tissue (Kilisch et al., 2023; Fridy et al., 2024).

How the size of these atypical MF synapses compares to that of MF synapses in the SL layer remains unclear. Electron microscopy 3D reconstruction studies estimate the average MF bouton’s cross-sectional area to be ∼5µm^2^ (Rollenhagen et al., 2007; Rollenhagen and Lübke, 2010; Murray et al., 2020), whereas our 2D average cluster size is ∼1.5µm^2^. This smaller size could be because our puncta only consist of the synaptic junction (colocalization of pre- and postsynaptic markers) portion of the larger MF structure. Future structural studies with high resolution imaging such as electron microscopy are needed to determine whether these atypical synapses are similar in size to the typical ‘MF’ synapses in the SL layer.

The reason why our cognitively enriched mice have fewer MF synapses remains an outstanding question. One possibility is increased synaptic pruning triggered by plasticity such as repeated bouts of long-term depression (LTD) (Bastrikova et al., 2008; Becker et al., 2008; Shinoda et al., 2010). The occurrence of LTD and LTP in the hippocampus largely depends on the nature of the behavioral learning task (Hagena and Manahan-Vaughan, 2024). At MF-CA3 synapses, exposure to a simple novel context facilitates LTP, while introducing large items and rearranging them, even when familiar, facilitates LTD (Kemp and Manahan-Vaughan, 2008) (Hagena and Manahan-Vaughan, 2011). Our ET includes large obstacles frequently changed and rearranged, suggesting LTP may occur initially, followed by ongoing LTD and pruning as mice continue learning in the same context.

The fact that MFs from adult-born DG neurons expand beyond the SL layer (Cole et al., 2020) raises the possibility that these atypical synapses preferentially belong to adult-born DG neurons. A recent study found that increases in the sparse activity of CA3 place cells due to enrichment requires neurogenesis (Ventura et al., 2024). EE can also rescue the survival of newborn neurons following ELS (Rule et al., 2021) and newborn neurons are more plastic (Ge et al., 2007; Massa et al., 2011). However, enrichment typically promotes growth factor signaling, enhancing DG-CA3 LTP and MF synaptogenesis (Gogolla et al., 2009; Bednarek and Caroni, 2011; Schildt et al., 2013; Cao et al., 2014). And, whether adult-born DG neurons have altered MF plasticity with CA3 remains unclear. Nevertheless, the proximity of atypical MF synapses to the spike initiation zone could tightly couple CA3 activity to changes in synaptic density. Alterations in the fraction of newborn neurons that form atypical MF synapses could represent a mechanism by which experience and neurogenesis fine tune hippocampal activity.

Our findings suggest a potential mechanism to restore spatial learning and hippocampal function in aged ELS mice. Studies show that age and ELS increase DG and CA3 excitability (Barnes and McNaughton, 1980; Wilson et al., 2005; Patrylo et al., 2007; Simkin et al., 2015; Villanueva-Castillo et al., 2017); changes that would increase population activity and impair the ability of the hippocampus to pattern separate (Fig. 5A) (Barnes et al., 1997; Tanila et al., 1997a, 1997b; Redish et al., 1998; Wilson et al., 2005; Jinde et al., 2012). Our cognitive enrichment causes a 27% decrease in the number of atypical MF synapses which could counteract high excitability by reducing the number of MF synapses. The resulting increase in sparse activity would favor pattern separation and improve spatial learning, as distinct contexts would have more orthogonal CA3 and CA1 representations (Fig. 5B) (McNaughton and Morris, 1987; McHugh et al., 2007).

Our results suggest that targeted cognitive enrichment training may be especially beneficial to individuals that have suffered from ELS. A potential therapeutic avenue is playing 3D action games, which significantly improve spatial reasoning and memory in humans. For instance, action video game players outperformed non-players on hippocampal-mediated memory functions such as mental rotation and spatial visualization tasks (Green and Bavelier, 2003; Uttal et al., 2013; Stark et al., 2015). Moreover, neuroimaging studies confirm that video game training increases gray matter in the hippocampus and prefrontal cortex (Kühn et al., 2014). Future studies, however, are needed to determine whether accessible interventions like 3D video games can mitigate the long-term cognitive and hippocampal structural deficits induced by early life adversity.

## MATERIALS AND METHODS

### Experimental animals

For the longitudinal study we used Thy1-GCaMP6f-GP5.17 (Jackson) mice due to the absence of reporter expression in the CA3. Upon reaching early-adulthood (2.5 months) GCaMP6f^+^ littermates were double-housed (after ELS) so that each cage had a mouse that ran ET and CT. This consisted of 4 cages of females (n=8) and 4 cages of males (n=8) for a total of (n=16). Due to the longevity of this study, 3 males and 1 female mouse died of natural causes before reaching the final 20-month timepoint. After removal of the GCaMP6f^+^ mice, there remained six male GCamP6f^-^littermates (3 cages). These were grouped with age typically reared (TR) age-matched C57BL/6J male mice purchased from Jackson (n=6) for the ELS vs TR experiment (Figure 2E & F).

### Early-life stress and cognitive enrichment

The experiments conducted were in accordance with the guidelines set by the Institutional Animal Care and Use Committee (IACUC) at the University of California, Irvine. We used the ELS protocol (developed in the Baram lab) whereby mouse litters receive limited bedding and nesting (LBN) conditions from postnatal days 2-10 in their rearing cages to modify maternal behavior during early development, (Fig. 1A, B). Briefly, on postnatal day 2, we replaced the normal bedding (∼6 litters) with a fitted metal grate (large enough to allow dropping to collect underneath) and two nestlets so that the dam could make a rudimentary nest. On postnatal day 10, we restored the normal bedding conditions. The control group (3 litters), referred to as typically reared (TR), remained in their normal cages. After reaching adulthood (3.5 months), we ran a subset of the ELS mice (N=16) on either enrichment (ET) or control track (CT) for ten weeks (3 x 30 mins sessions/week) (Fig. 1C), (Gattas et al., 2022). Briefly, the enrichment setup consisted of two juxtaposed square tracks: one containing obstacles, while the other had simple ramps with one-way doors located at diagonal corners. Initially, we trained both groups of mice (2 weeks) to run laps around the track loaded with simple ramps (12) while receiving a single milk reward dispensed from a mounted lick tube triggered at the conclusion of each lap. In the next phase, we introduced complex obstacles only to the ET group, which continued for 8 weeks (Fig 1B).

### Object location memory and novel object recognition

For each time point, we conducted a set of OLM (first week) and NOR (second week) behaviors over a two-week period (10 days total). The first three days of each week served as habituation sessions, during which animals explored empty square boxes (10 x 9 inches) for 10 minutes. On the training day (fourth day), mice were exposed to two identical objects for 10 minutes. On the test day (fifth day), one object was relocated to a new position (OLM, week 1, Fig. 2A, top) or replaced with a novel object in the same position (NOR, week 2, Fig. 2B, top). We used the same context box but different objects for each round of OLM and NOR to avoid object familiarity-related confounds. Boxes had unique markings on two of the walls (vertical and horizontal stripes) so that mice could easily associate the position of the objects with the box (Sup Fig. 2A and B). All behaviors were recorded with an overhead camera using infrared emitters for low-light conditions.

### Y-maze

The Y-maze setup had three identical arms (3.5 inches wide and 10.5 inches long) with transparent walls and was surrounded by distinct external cues (Fig. 2C). We always placed mice in the start arm at the beginning of each session. On training day (the first exposure to the Y-maze), we barricaded the to-be novel arm (opaque blocker) so that mice could only explore the two open arms (start and familiar) for 10 minutes. On test day (24 hrs later), we removed the blocker so that mice were free to explore all three arms for 5 mins. A single top-view camera captured training and testing session videos.

### Behavioral analysis

The behavior videos were analyzed manually using BORIS software (Friard and Gamba, 2016) to count the duration each animal spent within 2 cm of each object (OLM/ NOR) and moving inside the novel arm (spatial Y-maze). Memory performance in each behavior test was measured using discrimination ratio (DR), calculated as the ratio of time spent near novel conditions (object in a novel location for OLM or novel object for NOR or novel arm in spatial Y-maze) to time spent near familiar conditions (object in a familiar location for OLM or familiar object for NOR or familiar arm in spatial Y-maze).

### Immunofluorescence staining

Following isoflurane anesthesia, mice were transcardially perfused with cold phosphate buffer solution (PBS), followed by 4% paraformaldehyde (PFA), and the extracted brain was stored in 4% PFA at 4°C. Before slicing, brains were transferred to 30% sucrose/PBS solution and stored at 4°C for cryoprotection. Brains were sectioned at 40 μm using a cryostat (Thermo Scientific HM525 NX) at -20°C, and each section was stored in well plates containing cryoprotectant solution.

Anatomical targeting: Dorsal hippocampal sections were selected at approximately -1.8 mm from bregma (anterior-posterior coordinate) according to the Paxinos & Watson mouse brain atlas to visualize mossy fiber (MF) synapses in the distal CA3 region. This anatomical level was chosen to ensure regional consistency, as synaptogenesis is region-specific within hippocampal subfields, particularly in the septal versus temporal hippocampus.

Immunohistochemistry protocol: Initial attempts using traditional PSD-95 and presynaptic antibodies yielded poor results with low signal and poor colocalization. Subsequent optimization using nanobodies showed dramatic improvements. First, we incubated sections for 2 hours (at room temperature) in PBS blocking buffer containing 5% normal goat serum (NGS) and 0.3% Triton X-100. Sections were then incubated with nanobodies (purchased from NanoTag Biotechnologies) that contained the appropriate combination of either anti-PSD-95 nanobody (FluoTag®-X2 anti-PSD95, Cat No: N3702-AF647-L, 2 nM, Alexa647), anti-synaptotagmin 1 nanobody (FluoTag®-X2 anti-Synaptotagmin 1, Cat No: N2302-AF568-L, 0.2 nM, AZDye568) or VGLUT1 nanobody (FluoTag®-X2 anti-VGLUT1, Cat No: N1602-AF568-L, 1 nM, AZDye568). Incubation was performed with shaking at 4°C for 24 hours in PBS containing 0.3% Triton X-100. Sections were washed four times for 10 minutes each alternating between PBS and TBS. At these low concentrations, fluorescent signals were more prominent in the stratum pyramidale (SP) layer compared to the stratum lucidum (SL) layer. Sections were mounted onto slides with media containing DAPI (Invitrogen SlowFade Glass Soft-set Antifade Mountant with DAPI, catalogue #S36917).

### Gross morphological analysis

We took images of the entire dorsal hippocampus with a Keyence BZ-X1810 widefield fluorescence microscope with a 20x objective lens with DAPI and GCaMP filter cubes to visualize cell bodies and endogenous GCaMP6 signal. Images were loaded in Zeiss Zen software so that we could manually trace hippocampal subfields using the active contour tool. Following each outline, we took reported areas and normalized them to the total hippocampal area. To determine number of CA3 neuron we counted the number of easily identifiable large dark circles in the tiled images (described below). All tracing and counting was done by a double-blind observer.

### Mossy fiber synaptic analysis

Confocal imaging: Tiled images of the distal CA3 bend were captured using an LSM 900 microscope equipped with Airyscan 2 and a 63X objective lens. PSD-95 and synaptotagmin-1 were excited using 653 nm and 568 nm diode lasers, respectively. Images were over-sampled (2x) to facilitate super-resolution post-processing with Zeiss’ Super Resolution Airyscan mode (2D, auto). No additional deconvolution was applied beyond the Airyscan super-resolution processing. Images were manually aligned using Imaris Stitcher, creating two composite panels per mouse (approximately 20 stitched images for each z-plane panel) selected systematically from the dorsal and ventral aspects of the CA3 bend. These panels were roughly 8,000 x 10,000 pixels (∼350 x 420 µm), 16-bit TIFF files, featuring red (PSD-95) and green (synaptotagmin-1) channels. Imaging parameters including laser power (2.3% for 640 nm and 3.0% for 561 nm), gain, and offset were kept constant across all samples to ensure quantitative comparisons. As expected, nanobody penetration was lowest in the middle of the slice so for each mouse we constructed two composite panels at 10 μm and 30 μm positions within the 40 μm slice. We used the same settings for non-tiled images with the following additions: laser 405 nm at 1.5% and laser 488 nm power at 1.3%. Z-stacks consisted of 15 optical sections centered at 10 μm with a 1 μm step size. Reconstructed 3D images of synapses surrounding the cell body are max Z projections of all 15 slices.

Quantitative analysis: For subsequent analysis on the tiled images, FIJI was used to crop images focusing on the pyramidal cell bodies of the SP layer. These cropped, composite images served as input for the MF synapse detection algorithm. Each channel was independently thresholded based on fluorescence intensity to create binary masks of equal size. Threshold values of 515 for PSD-95 and 350 for Synaptotagmin were selected because they captured all identifiable clusters while excluding apparent noise, with similar mask sizes observed with threshold variations within a 5% range. The masks were combined using a pixel-wise AND operation and applied to the original 2-channel image, retaining only pixel values above the respective thresholds, with all other pixels set to 0 (black). Clusters smaller than 150 pixels were removed from subsequent analysis to eliminate imaging artifacts, so only clusters with an area exceeding 75% of 0.37 μm² (0.27 μm²) were included. Large images were processed in parallel using smaller blocks of 500 x 500 pixels, with a flood-fill algorithm applied to every fourth pixel (provided it was a non-zero value) to detect cluster masks. Visual inspections were performed on two images by manual counting to validate the automated detection, showing 97% agreement between automated and manual counts. Duplicate detections at block edges were filtered to ensure a unique set of clusters. Mossy fiber bouton density was quantified as the number of synaptic clusters per CA3 neuron within the pyramidal cell layer, normalized across the two composite panels per mouse. All image acquisition and quantitative analyses were performed by investigators blinded to experimental groups. The analysis code is freely available online at GitHub [https://github.com/bainro/MF-synapse-analysis].

### Statistical Analysis

Behavioral tests were analyzed using repeated measures and two-way ANOVA designs (α = 0.05). Object location memory (OLM) and novel object recognition (NOR) were analyzed using three-way repeated measures ANOVA (Age × Group × Sex) with n=16 mice total (ELS+ET: 4M and 4F, n=8; ELS+CT: 4M and 4F, n=8). For OLM, only the Group main effect was statistically significant [F(1,12) = 8.80, p = 0.012], while all other main effects and interactions were non-significant: Age [F(1,12) = 1.69, p = 0.218], Sex [F(1,12) = 0.97, p = 0.345], Age × Group [F(1,12) = 0.16, p = 0.700], Age × Sex [F(1,12) = 0.081, p = 0.780], Group × Sex [F(1,12) = 0.0004, p = 0.984], and Age × Group × Sex [F(1,12) = 0.33, p = 0.579]. For NOR, no main effects or interactions reached statistical significance: Age [F(1,12) = 2.93, p = 0.113], Group [F(1,12) = 0.074, p = 0.791], Sex [F(1,12) = 0.11, p = 0.744], Age × Group [F(1,12) = 1.66, p = 0.222], Age × Sex [F(1,12) = 0.53, p = 0.479], Group × Sex [F(1,12) = 4.48, p = 0.056], and Age × Group × Sex [F(1,12) = 0.55, p = 0.472]. Object exploration time (OLM & NOR combined) was analyzed using a two-way ANOVA (Test Type × Group) with the same sample sizes. Only the Test Type main effect was statistically significant [F(2,88) = 30.86, p < 0.0001], while Group [F(1,88) = 0.886, p = 0.349] and Test Type × Group interaction [F(2,88) = 0.101, p = 0.904] were non-significant. Y-maze data were analyzed using two-way ANOVA (Group × Sex) with n=12 mice total (ELS+ET: 2M and 3F, n=5; ELS+CT: 3M and 4F, n=7). Only the Group main effect was statistically significant [F(1,8) = 19.89, p = 0.002], while Sex [F(1,8) = 1.81, p = 0.215] and Group × Sex interaction [F(1,8) = 0.81, p = 0.393] were non-significant.

Synaptic analyses were conducted using two-way ANOVA (Group × Sex, α = 0.05) with n=12 mice total (ELS+ET: 2M and 3F, n=5; ELS+CT: 3M and 4F, n=7). A significant Group main effect was found for cluster count [F(1,8) = 6.07, p = 0.039], with Sex [F(1,8) = 0.043, p = 0.841] and Group × Sex interaction [F(1,8) = 0.073, p = 0.794] being non-significant. For cluster size, no main effects or interactions reached statistical significance: Group [F(1,8) = 4.36, p = 0.070], Sex [F(1,8) = 0.0009, p = 0.976], and Group × Sex interaction [F(1,8) = 0.42, p = 0.535].

Gross morphological analyses were conducted using two-way ANOVA (Group × Sex, α = 0.05) with n=12 mice total (ELS+ET: 2M and 3F, n=5; ELS+CT: 3M and 4F, n=7). All hippocampal morphology measures showed non-significant results including CA3 neuron density [Group: F(1,8) = 1.734, p = 0.224; Sex: F(1,8) = 0.026, p = 0.876; Group × Sex: F(1,8) = 0.012, p = 0.917], hilus area [Group: F(1,8) = 0.318, p = 0.589; Sex: F(1,8) = 0.268, p = 0.619; Group × Sex: F(1,8) = 0.117, p = 0.741], mossy fiber area [Group: F(1,8) = 0.596, p = 0.462; Sex: F(1,8) = 4.265, p = 0.073; Group × Sex: F(1,8) = 0.144, p = 0.714], and dentate gyrus area [Group: F(1,8) = 0.024, p = 0.881; Sex: F(1,8) = 4.056, p = 0.079; Group × Sex: F(1,8) = 0.098, p = 0.762]. Sex effects approached significance for both mossy fiber area (p = 0.073) and dentate gyrus area (p = 0.079).

Summary of significant findings: Cognitive enrichment training (ET) significantly improved spatial memory performance in ELS mice, as evidenced by significant Group effects in both OLM (p = 0.012) and Y-maze (p = 0.002) tasks, but not in the hippocampus-independent NOR task. At the synaptic level, ET training significantly reduced the number of atypical mossy fiber synaptic clusters surrounding CA3 cell bodies (p = 0.039), while gross hippocampal morphology remained unchanged. These findings suggest that cognitive enrichment rescues ELS-induced spatial memory deficits through selective modifications of synaptic connectivity rather than gross structural changes.

## AUTHOR’S CONTRIBUTIONS

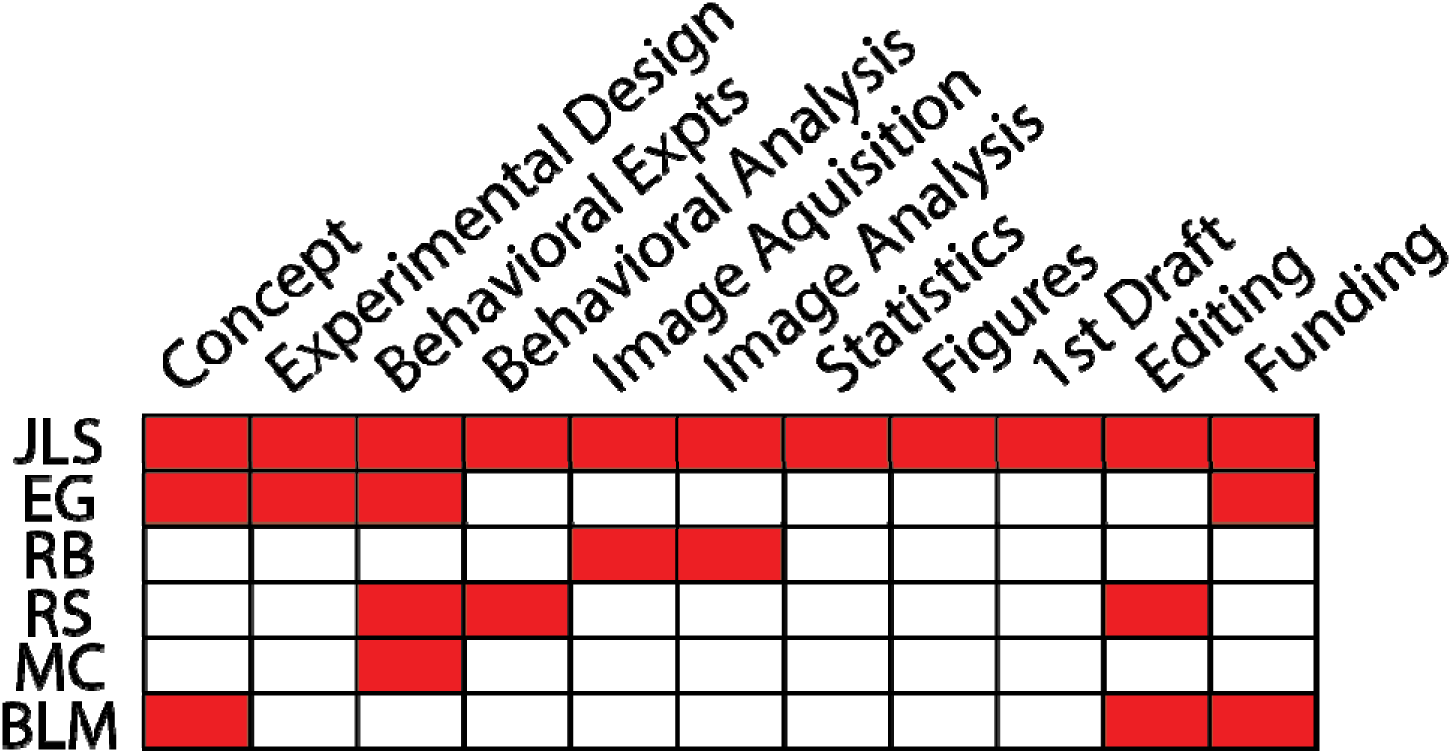

## ACKNOWLEDGMENTS

We thank Tallie Z. Baram MD, PhD for her advice on experimental design and members of her lab, Annabel Short, PhD and Racheal Hokenson, PhD who provided us with details about conducting and analyzing the ELS and behavioral experiments. In the McNaughton lab, we thank Mariya Vodyanyk for establishing behavioral analysis methods as well as Aida Andujo and Varleen Kaur for excellent technical assistance with mouse behavior. This work was initiated by a 1-year UCI conte seed grant to JLS and EG and further supported by NIH BRAIN grants: R01 NS121764 and RF1 NS132041 to BLM. This study was made possible in part through access to the Optical Biology Core Facility of the Developmental Biology Center, a shared resource supported by the Cancer Center Support Grant (CA-62203) and NIH-S10OD032327-01.

## LITERATURE CITED

Adams MM, Donohue HS, Linville MC, Iversen EA, Newton IG, Brunso-Bechtold JK (2010) Age-related synapse loss in hippocampal CA3 is not reversed by caloric restriction. Neuroscience 171:373–382.

Almas AN, Degnan KA, Nelson CA, Zeanah CH, Fox NA (2016) IQ at Age 12 Following a History of Institutional Care: Findings From the Bucharest Early Intervention Project. Dev Psychol 52:1858–1866.

Alwis DS, Rajan R (2014) Environmental enrichment and the sensory brain: the role of enrichment in remediating brain injury. Front Syst Neurosci 8:156.

Amaral DG, Dent JA (1981) Development of the mossy fibers of the dentate gyrus: I. A light and electron microscopic study of the mossy fibers and their expansions. J Comp Neurol 195:51– 86.

Artola A, von Frijtag JC, Fermont PC, Gispen WH, Schrama LH, Kamal A, Spruijt BM. Long-lasting modulation of the induction of LTD and LTP in rat hippocampal CA1 by behavioural stress and environmental enrichment. Eur J Neurosci. 2006 Jan;23(1):261–72. doi: 10.1111/j.1460-9568.2005.04552.x. PMID: 16420435.

Banks E, Gutekunst C-A, Vargish GA, Eaton A, Pelkey KA, McBain CJ, Zheng JQ, Oláh VJ, Rowan MJM (2024) An enhancer-AAV approach selectively targeting dentate granule cells of the mouse hippocampus. Cell Rep Methods 4:100684.

Barker GRI, Warburton EC (2011) When Is the Hippocampus Involved in Recognition Memory? J Neurosci 31:10721–10731.

Barnes C, McNaughton B (1980) Physiological compensation for loss of afferent synapses in rat hippocampal granule cells during senescence. J Physiology 309:473–485.

Barnes CA, Suster MS, Shen J, McNaughton BL (1997) Multistability of cognitive maps in the hippocampus of old rats. Nature 388:272–275.

Bastrikova N, Gardner GA, Reece JM, Jeromin A, Dudek SM (2008) Synapse elimination accompanies functional plasticity in hippocampal neurons. Proc Natl Acad Sci 105:3123– 3127.

Becker N, Wierenga CJ, Fonseca R, Bonhoeffer T, Nägerl UV (2008) LTD Induction Causes Morphological Changes of Presynaptic Boutons and Reduces Their Contacts with Spines. Neuron 60:590–597.

Beckett C, Maughan B, Rutter M, Castle J, Colvert E, Groothues C, Hawkins A, Kreppner J, O’Connor TG, Stevens S, Sonuga-Barke EJS (2007) Scholastic Attainment Following Severe Early Institutional Deprivation: A Study of Children Adopted from Romania. J Abnorm Child Psychol 35:1063–1073.

Bednarek E, Caroni P (2011) β-Adducin Is Required for Stable Assembly of New Synapses and Improved Memory upon Environmental Enrichment. Neuron 69:1132–1146.

Bennett JC, McRae PA, Levy LJ, Frick KM. Long-term continuous, but not daily, environmental enrichment reduces spatial memory decline in aged male mice. Neurobiol Learn Mem. 2006 Mar;85(2):139–52. doi: 10.1016/j.nlm.2005.09.003. Epub 2005 Oct 26. PMID: 16256380.

Bilkey DK, Cheyne KR, Eckert MJ, Lu X, Chowdhury S, Worley PF, Crandall JE, Abraham WC (2017) Exposure to complex environments results in more sparse representations of space in the hippocampus. Hippocampus 27:1178–1191.

Bolton JL, Schulmann A, Garcia-Curran MM, Regev L, Chen Y, Kamei N, Shao M, Singh-Taylor A, Jiang S, Noam Y, Molet J, Mortazavi A, Baram TZ. Unexpected Transcriptional Programs Contribute to Hippocampal Memory Deficits and Neuronal Stunting after Early-Life Adversity. Cell Rep. 2020 Dec 15;33(11):108511.

Bos KJ, Fox N, Zeanah CH, III CAN (2009) Effects of Early Psychosocial Deprivation on the Development of Memory and Executive Function. Front Behav Neurosci 3:16.

Bramati G, Stauffer P, Nigri M, Wolfer DP, Amrein I (2023) Environmental enrichment improves hippocampus-dependent spatial learning in female C57BL/6 mice in novel IntelliCage sweet reward-based behavioral tests. Front Behav Neurosci 17:1256744.

Brunson KL, Eghbal-Ahmadi M, Bender R, Chen Y, Baram TZ (2001) Long-term, progressive hippocampal cell loss and dysfunction induced by early-life administration of corticotropin-releasing hormone reproduce the effects of early-life stress. Proc Natl Acad Sci 98:8856– 8861.

Brunson KL, Kramár E, Lin B, Chen Y, Colgin LL, Yanagihara TK, Lynch G, Baram TZ (2005) Mechanisms of Late-Onset Cognitive Decline after Early-Life Stress. J Neurosci 25:9328– 9338.

Cai X, Bai X, Zhou S (2024) Childhood adversities and memory function in later life: the mediating role of activity participation. BMC Geriatr 24:536.

Cao W, Duan J, Wang X, Zhong X, Hu Z, Huang F, Wang H, Zhang J, Li F, Zhang J, Luo X, Li C-Q (2014) Early enriched environment induces an increased conversion of proBDNF to BDNF in the adult rat’s hippocampus. Behav Brain Res 265:76–83.

Cabezas C, Irinopoulou T, Gauvain G, Poncer JC (2012) Presynaptic but not postsynaptic GABA signaling at unitary mossy fiber synapses. J Neurosci 32:11835–11840.

Carasatorre M, Ochoa-Alvarez A, Velázquez-Campos G, Lozano-Flores C, Díaz-Cintra SY, Ramírez-Amaya V (2015) Hippocampal Synaptic Expansion Induced by Spatial Experience in Rats Correlates with Improved Information Processing in the Hippocampus. PLoS ONE 10:e0132676.

Chen Y, Molet J, Lauterborn JC, Trieu BH, Bolton JL, Patterson KP, Gall CM, Lynch G, Baram TZ (2016) Converging, Synergistic Actions of Multiple Stress Hormones Mediate Enduring Memory Impairments after Acute Simultaneous Stresses. J Neurosci 36:11295–11307.

Cheng ST, Liu S, Ou-Yang B, Dai XY, Cheng L. Specific Effects of Characteristics of Enriched Environment on Innovative Problem Solving by Animals. Psychol Sci. 2022 Jul;33(7):1097–1111. doi: 10.1177/09567976211070562. Epub 2022 Jul 1. PMID: 35776087.

Cohen SJ, Jr. RW (2015) Assessing rodent hippocampal involvement in the novel object recognition task. A review. Behavioural Brain Research 285:105–117.

Cole JD, Espinueva DF, Seib DR, Ash AM, Cooke MB, Cahill SP, O’Leary TP, Kwan SS, Snyder JS (2020) Adult-Born Hippocampal Neurons Undergo Extended Development and Are Morphologically Distinct from Neonatally-Born Neurons. J Neurosci 40:5740–5756.

Connor JR, Wang EC, Diamond MC. Increased length of terminal dendritic segments in old adult rats’ somatosensory cortex: an environmentally induced response. Exp Neurol. 1982 Nov;78(2):466–70. doi: 10.1016/0014-4886(82)90064-4. PMID:

Cortese GP, Olin A, O’Riordan K, Hullinger R, Burger C. Environmental enrichment improves hippocampal function in aged rats by enhancing learning and memory, LTP, and mGluR5-Homer1c activity. Neurobiol Aging. 2018 Mar;63:1–11. doi: 10.1016/j.neurobiolaging.2017.11.004. Epub 2017 Nov 16. PMID: 29207276; PMCID: PMC5801151.

Cui M, Yang Y, Yang J, Zhang J, Han H, Ma W, Li H, Mao R, Xu L, Hao W, Cao J (2006) Enriched environment experience overcomes the memory deficits and depressive-like behavior induced by early life stress. Neurosci Lett 404:208–212.

Dailey M, Buchanan J, Bergles D, Smith S (1994) Mossy fiber growth and synaptogenesis in rat hippocampal slices in vitro. J Neurosci 14:1060–1078.

Ding R, He P (2021) Associations between childhood adversities and late-life cognitive function: Potential mechanisms. Soc Sci Med 291:114478.

Eckenrode J, Laird M, Doris J (1993) School Performance and Disciplinary Problems Among Abused and Neglected Children. Dev Psychol 29:53–62.

Eigsti I-M, Weitzman C, Schuh J, Marchena A de, Casey BJ (2011) Language and cognitive outcomes in internationally adopted children. Dev Psychopathol 23:629–646.

Engineer ND, Percaccio CR, Pandya PK, Moucha R, Rathbun DL, Kilgard MP. Environmental enrichment improves response strength, threshold, selectivity, and latency of auditory cortex neurons. J Neurophysiol. 2004 Jul;92(1):73–82.

Fenoglio KA, Brunson KL, Baram TZ (2006) Hippocampal neuroplasticity induced by early-life stress: Functional and molecular aspects. Front Neuroendocrin 27:180–192.

Freund TF, Buzsáki G (1996) Interneurons of the hippocampus. Hippocampus 6:347–470.

Friard O, Gamba M (2016) BORIS: a free, versatile open-source event-logging software for video/audio coding and live observations. Methods Ecol Evol 7:1325–1330.

Fridy PC, Farrell RJ, Molloy KR, Keegan S, Wang J, Jacobs EY, Li Y, Trivedi J, Sehgal V, Fenyö D, Wu Z, Chait BT, Rout MP (2024) A new generation of nanobody research tools using improved mass spectrometry-based discovery methods. J Biol Chem 300:107623.

Galimberti I, Gogolla N, Alberi S, Santos AF, Muller D, Caroni P (2006) Long-Term Rearrangements of Hippocampal Mossy Fiber Terminal Connectivity in the Adult Regulated by Experience. Neuron 50:749–763.

Gattas S, Collett HA, Huff AE, Creighton SD, Weber SE, Buckhalter SS, Manning SA, Ryait HS, McNaughton BL, Winters BD (2022) A rodent obstacle course procedure controls delivery of enrichment and enhances complex cognitive functions. Npj Sci Learn 7:21.

Ge S, Yang C, Hsu K, Ming G, Song H (2007) A Critical Period for Enhanced Synaptic Plasticity in Newly Generated Neurons of the Adult Brain. Neuron 54:559–566.

Gogolla N, Galimberti I, Deguchi Y, Caroni P (2009) Wnt Signaling Mediates Experience-Related Regulation of Synapse Numbers and Mossy Fiber Connectivities in the Adult Hippocampus. Neuron 62:510–525.

Green CS, Bavelier D (2003) Action video game modifies visual selective attention. Nature 423:534–537.

Gutiérrez R, Heinemann U, Gaiarsa JL, Romo-Parra H (2003) Plasticity of the GABAergic phenotype of the “glutamatergic” granule cells of the rat dentate gyrus. J Neurosci 23:5594– 5598.

Hagena H, Manahan-Vaughan D (2011) Learning-Facilitated Synaptic Plasticity at CA3 Mossy Fiber and Commissural–Associational Synapses Reveals Different Roles in Information Processing. Cereb Cortex 21:2442–2449.

Hagena H, Manahan-Vaughan D (2024) Interplay of hippocampal long-term potentiation and long-term depression in enabling memory representations. Philos Trans B 379:20230229.

Henze DA, Urban NN, Barrionuevo G (2000) The multifarious hippocampal mossy fiber pathway: a review. Neuroscience 98:407–427.

Hoeijmakers L, Amelianchik A, Verhaag F, Kotah J, Lucassen PJ, Korosi A (2018) Early-Life Stress Does Not Aggravate Spatial Memory or the Process of Hippocampal Neurogenesis in Adult and Middle-Aged APP/PS1 Mice. Front Aging Neurosci 10:61.

Humphreys KL, King LS, Sacchet MD, Camacho MC, Colich NL, Ordaz SJ, Ho TC, Gotlib IH (2019) Evidence for a sensitive period in the effects of early life stress on hippocampal volume. Dev Sci 22:e12775.

Ivy AS, Rex CS, Chen Y, Dube C, Maras PM, Grigoriadis DE, Gall CM, Lynch G, Baram TZ (2010) Hippocampal Dysfunction and Cognitive Impairments Provoked by Chronic Early-Life Stress Involve Excessive Activation of CRH Receptors. J Neurosci 30:13005–13015.

Jinde S, Zsiros V, Jiang Z, Nakao K, Pickel J, Kohno K, Belforte JE, Nakazawa K (2012) Hilar Mossy Cell Degeneration Causes Transient Dentate Granule Cell Hyperexcitability and Impaired Pattern Separation. Neuron 76:1189–1200.

Jung CK, Herms J. Structural dynamics of dendritic spines are influenced by an environmental enrichment: an in vivo imaging study. Cereb Cortex. 2014 Feb;24(2):377–84. doi: 10.1093/cercor/bhs317. Epub 2012 Oct 18. PMID: 23081882.

Kawamoto M, Takagishi H, Ishihara T, Takagi S, Kanai R, Sugihara G, Takahashi H, Matsuda T (2023) Hippocampal volume mediates the relationship of parental rejection in childhood with social cognition in healthy adults. Sci Rep 13:19167.

Kemp A, Manahan-Vaughan D (2008) The Hippocampal CA1 Region and Dentate Gyrus Differentiate between Environmental and Spatial Feature Encoding through Long-Term Depression. Cereb Cortex 18:968–977.

Kempermann G (2019) Environmental enrichment, new neurons and the neurobiology of individuality. Nat Rev Neurosci 20:235–245.

Kilisch M, Gere-Becker M, Wüstefeld L, Bonnas C, Crauel A, Mechmershausen M, Martens H, Götzke H, Opazo F, Frey S (2023) Simple and Highly Efficient Detection of PSD95 Using a Nanobody and Its Recombinant Heavy-Chain Antibody Derivatives. Int J Mol Sci 24:7294.

Klausberger T, Somogyi P (2008) Neuronal diversity and temporal dynamics: the unity of hippocampal circuit operations. Science 321:53–57.

Kloc ML, Velasquez F, Niedecker RW, Barry JM, Holmes GL. Disruption of hippocampal rhythms via optogenetic stimulation during the critical period for memory development impairs spatial cognition. Brain Stimul. 2020 Nov-Dec;13(6):1535–1547.

Koyama Y, Fujiwara T, Murayama H, Machida M, Inoue S, Shobugawa Y (2022) Association between adverse childhood experiences and brain volumes among Japanese community-dwelling older people: Findings from the NEIGE study. Child Abus Negl 124:105456.

Kühn S, Gleich T, Lorenz RC, Lindenberger U, Gallinat J (2014) Playing Super Mario induces structural brain plasticity: gray matter changes resulting from training with a commercial video game. Mol Psychiatry 19:265–271.

Leinekugel X, Khazipov R, Cannon R, Hirase H, Ben-Ari Y, Buzsáki G (2002) Correlated Bursts of Activity in the Neonatal Hippocampus in Vivo. Science 296:2049–2052.

LeMessurier AM, Laboy-Juárez KJ, McClain K, Chen S, Nguyen T, Feldman DE. Enrichment drives emergence of functional columns and improves sensory coding in the whisker map in L2/3 of mouse S1. Elife. 2019 Aug 16;8:e46321. doi: 10.7554/eLife.46321. PMID: 31418693; PMCID: PMC6697414.

Ma J, Yang Y, Wan Y, Shen C, Qiu P (2021) The influence of childhood adversities on mid to late cognitive function: From the perspective of life course. PLoS ONE 16:e0256297.

Malave L, Dijk MT van, Anacker C (2022) Early life adversity shapes neural circuit function during sensitive postnatal developmental periods. Transl Psychiat 12:306.

Manno FAM, Kumar R, An Z, Khan MS, Su J, Liu J, Wu EX, He J, Feng Y, Lau C (2022) Structural and Functional Hippocampal Correlations in Environmental Enrichment During the Adolescent to Adulthood Transition in Mice. Front Syst Neurosci 15:807297.

Massa F, Koehl M, Koelh M, Wiesner T, Grosjean N, Revest J-M, Piazza P-V, Abrous DN, Oliet SHR (2011) Conditional reduction of adult neurogenesis impairs bidirectional hippocampal synaptic plasticity. Proc Natl Acad Sci 108:6644–6649.

McHugh TJ, Jones MW, Quinn JJ, Balthasar N, Coppari R, Elmquist JK, Lowell BB, Fanselow MS, Wilson MA, Tonegawa S (2007) Dentate Gyrus NMDA Receptors Mediate Rapid Pattern Separation in the Hippocampal Network. Science 317:94–99.

McNaughton BL, Morris RGM (1987) Hippocampal synaptic enhancement and information storage within a distributed memory system. Trends Neurosci 10:408–415.

Molet J, Heins K, Zhuo X, Mei YT, Regev L, Baram TZ, Stern H (2016a) Fragmentation and high entropy of neonatal experience predict adolescent emotional outcome. Transl Psychiat 6:e702–e702.

Molet J, Maras PM, Kinney-Lang E, Harris NG, Rashid F, Ivy AS, Solodkin A, Obenaus A, Baram TZ (2016b) MRI uncovers disrupted hippocampal microstructure that underlies memory impairments after early-life adversity. Hippocampus 26:1618–1632.

Murray KD, Liu X-B, King AN, Luu JD, Cheng H-J (2020) Age-Related Changes in Synaptic Plasticity Associated with Mossy Fiber Terminal Integration during Adult Neurogenesis. eNeuro 7:ENEURO.0030-20.2020.

Naninck EFG, Hoeijmakers L, Kakava-Georgiadou N, Meesters A, Lazic SE, Lucassen PJ, Korosi A (2015) Chronic early life stress alters developmental and adult neurogenesis and impairs cognitive function in mice. Hippocampus 25:309–328.

Naninck EFG, Oosterink JE, Yam K, Vries LP, Schierbeek H, Goudoever JB, Verkaik-Schakel R, Plantinga JA, Plosch T, Lucassen PJ, Korosi A (2017) Early micronutrient supplementation protects against early stress-induced cognitive impairments. Faseb J 31:505–518.

Nelson CA, Zeanah CH, Fox NA, Marshall PJ, Smyke AT, Guthrie D (2007) Cognitive Recovery in Socially Deprived Young Children: The Bucharest Early Intervention Project.

Oliveira AMM, Hawk JD, Abel T, Havekes R (2010) Post-training reversible inactivation of the hippocampus enhances novel object recognition memory. Learn Mem 17:155–160.

Patrylo PR, Tyagi I, Willingham AL, Lee S, Williamson A (2007) Dentate Filter Function Is Altered in a Proepileptic Fashion during Aging. Epilepsia 48:1964–1978.

Pintori N, Piva A, Mottarlini F, Díaz FC, Maggi C, Caffino L, Fumagalli F, Chiamulera C. Brief exposure to enriched environment rapidly shapes the glutamate synapses in the rat brain: A metaplastic fingerprint. Eur J Neurosci. 2024 Mar;59(5):982–995. doi: 10.1111/ejn.16279. Epub 2024 Feb 20. PMID: 38378276.

Pollak SD, Nelson CA, Schlaak MF, Roeber BJ, Wewerka SS, Wiik KL, Frenn KA, Loman MM, Gunnar MR (2010) Neurodevelopmental Effects of Early Deprivation in Postinstitutionalized Children. Child Dev 81:224–236.

Qin L, Marrs GS, McKim R, Dailey ME (2001) Hippocampal mossy fibers induce assembly and clustering of PSD95-containing postsynaptic densities independent of glutamate receptor activation. J Comp Neurol 440:284–298.

Ramı rez-Amaya V, Balderas I, Sandoval J, Escobar ML, Bermúdez-Rattoni F (2001) Spatial Long-Term Memory Is Related to Mossy Fiber Synaptogenesis. J Neurosci 21:7340–7348.

Redish AD, McNaughton BL, Barnes CA (1998) Reconciling Barnes et al. (1997) and Tanila et al. (1997a,b). Hippocampus 8:438–443.

Rice CJ, Sandman CA, Lenjavi MR, Baram TZ (2008) A Novel Mouse Model for Acute and Long-Lasting Consequences of Early Life Stress. Endocrinology 149:4892–4900.

Rocha M, Wang D, Avila-Quintero V, Bloch MH, Kaffman A. Deficits in hippocampal-dependent memory across different rodent models of early life stress: systematic review and meta-analysis. Transl Psychiatry. 2021 Apr 20;11(1):231.

Rollenhagen A, Lübke JHR (2010) The Mossy Fiber Bouton: the “Common” or the “Unique” Synapse? Front Synaptic Neurosci 2:2.

Rollenhagen A, Sätzler K, Rodríguez EP, Jonas P, Frotscher M, Lübke JHR (2007) Structural Determinants of Transmission at Large Hippocampal Mossy Fiber Synapses. J Neurosci 27:10434–10444.

Rountree-Harrison D, Burton TJ, Leamey CA, Sawatari A. Environmental Enrichment Expedites Acquisition and Improves Flexibility on a Temporal Sequencing Task in Mice. Front Behav Neurosci. 2018 Mar 15;12:51. doi: 10.3389/fnbeh.2018.00051. PMID: 29599712; PMCID: PMC5862792.

Rule L, Yang J, Watkin H, Hall J, Brydges NM (2021) Environmental enrichment rescues survival and function of adult-born neurons following early life stress. Mol Psychiatry 26:1898–1908.

Schildt S, Endres T, Lessmann V, Edelmann E (2013) Acute and chronic interference with BDNF/TrkB-signaling impair LTP selectively at mossy fiber synapses in the CA3 region of mouse hippocampus. Neuropharmacology 71:247–254.

Scholz J, Allemang-Grand R, Dazai J, Lerch JP (2015) Environmental enrichment is associated with rapid volumetric brain changes in adult mice. NeuroImage 109:190–198.

Schwegler H, Crusio WE, Brust I (1990) Hippocampal mossy fibers and radial-maze learning in the mouse: A correlation with spatial working memory but not with non-spatial reference memory. Neuroscience 34:293–298.

Shinoda Y, Tanaka T, Tominaga-Yoshino K, Ogura A (2010) Persistent Synapse Loss Induced by Repetitive LTD in Developing Rat Hippocampal Neurons. PLoS ONE 5:e10390.

Short AK, Maras PM, Pham AL, Ivy AS, Baram TZ. Blocking CRH receptors in adults mitigates age-related memory impairments provoked by early-life adversity. Neuropsychopharmacology. 2020 Feb;45(3):515–523.

Simkin D, Hattori S, Ybarra N, Musial TF, Buss EW, Richter H, Oh MM, Nicholson DA, Disterhoft JF (2015) Aging-Related Hyperexcitability in CA3 Pyramidal Neurons Is Mediated by Enhanced A-Type K+ Channel Function and Expression. J Neurosci 35:13206–13218.

Speisman RB, Kumar A, Rani A, Pastoriza JM, Severance JE, Foster TC, Ormerod BK. Environmental enrichment restores neurogenesis and rapid acquisition in aged rats. Neurobiol Aging. 2013 Jan;34(1):263–74. doi: 10.1016/j.neurobiolaging.2012.05.023. Epub 2012 Jul 12. PMID: 22795793; PMCID: PMC3480541.

Spratt EG, Friedenberg SL, Swenson CC, Larosa A, Bellis MDD, Macias MM, Summer AP, Hulsey TC, Runyan DK, Brady KT (2012) The Effects of Early Neglect on Cognitive, Language, and Behavioral Functioning in Childhood. Psychology 03:175–182.

Stark CE, Clemenson GD, Aluru K, Hatamian N (2015) Playing three-dimensional video games can improve hippocampus-associated memory discriminability. J Neurosci 35:16116–16125.

Tanila H, Shapiro M, Gallagher M, Eichenbaum H (1997a) Brain Aging: Changes in the Nature of Information Coding by the Hippocampus. J Neurosci 17:5155–5166.

Tanila H, Shapiro ML, Eichenbaum H (1997b) Discordance of spatial representation in ensembles of hippocampal place cells. Hippocampus 7:613–623.

Teicher MH, Samson JA, Anderson CM, Ohashi K (2016) The effects of childhood maltreatment on brain structure, function and connectivity. Nat Rev Neurosci 17:652–666.

Tizard B, Rees J (1974) A comparison of the effects of adoption, restoration to the natural mother, and continued institutionalization on the cognitive development of four-year-old children. Child Dev 45:92–99.

Toscano-Silva M, Silva SG da, Scorza FA, Bonvent JJ, Cavalheiro EA, Arida RM (2010) Hippocampal mossy fiber sprouting induced by forced and voluntary physical exercise. Physiol Behav 101:302–308.

Uchigashima M, Fukaya M, Watanabe M, Kamiya H (2007) Evidence against GABA release from glutamatergic mossy fiber terminals in the developing hippocampus. J Neurosci 27:8088–8100.

Urban NN, Henze DA, Barrionuevo G (2001) Revisiting the role of the hippocampal mossy fiber synapse. Hippocampus 11:408–417.

Uttal DH, Meadow NG, Tipton E, Hand LL, Alden AR, Warren C, Newcombe NS (2013) The malleability of spatial skills: a meta-analysis of training studies. Psychol Bull 139:352–402.

Ventura S, Duncan S, Ainge JA (2024) Increased flexibility of CA3 memory representations following environmental enrichment. Curr Biol 34:2011–2019.e7.

Villanueva-Castillo C, Tecuatl C, Herrera-López G, Galván EJ (2017) Aging-related impairments of hippocampal mossy fibers synapses on CA3 pyramidal cells. Neurobiol Aging 49:119– 137.

Walker C-D, Bath KG, Joels M, Korosi A, Larauche M, Lucassen PJ, Morris MJ, Raineki C, Roth TL, Sullivan RM, Taché Y, Baram TZ (2017) Chronic early life stress induced by limited bedding and nesting (LBN) material in rodents: critical considerations of methodology, outcomes and translational potential. Ann Ny Acad Sci 20:421–448.

Walker MC, Ruiz A, Kullmann DM (2001) Monosynaptic GABAergic signaling from dentate to CA3 with a pharmacological and physiological profile typical of mossy fiber synapses. Neuron 29:703–715.

Wang X-D, Rammes G, Kraev I, Wolf M, Liebl C, Scharf SH, Rice CJ, Wurst W, Holsboer F, Deussing JM, Baram TZ, Stewart MG, Müller MB, Schmidt MV (2011) Forebrain CRF1 Modulates Early-Life Stress-Programmed Cognitive Deficits. J Neurosci 31:13625–13634.

Williams BM, Luo Y, Ward C, Redd K, Gibson R, Kuczaj SA, McCoy JG. Environmental enrichment: effects on spatial memory and hippocampal CREB immunoreactivity. Physiol Behav. 2001 Jul;73(4):649–58. doi: 10.1016/s0031-9384(01)00543-1. PMID: 11495671.

Wilson IA, Ikonen S, Gallagher M, Eichenbaum H, Tanila H (2005) Age-Associated Alterations of Hippocampal Place Cells Are Subregion Specific. J Neurosci 25:6877–6886.

Wodarski JS, Kurtz PD, Gaudin JM, Howing PT (1990) Maltreatment and the School-Age Child: Major Academic, Socioemotional, and Adaptive Outcomes. Soc Work 35:506–513.

Woodcock EA, Richardson R. Effects of environmental enrichment on rate of contextual processing and discriminative ability in adult rats. Neurobiol Learn Mem. 2000 Jan;73(1):1–10. doi: 10.1006/nlme.1999.3911. PMID: 10686119.

Youssef M, Atsak P, Cardenas J, Kosmidis S, Leonardo ED, Dranovsky A (2019) Early life stress delays hippocampal development and diminishes the adult stem cell pool in mice. Sci Rep 9:4120.

Zeleznikow-Johnston A, Burrows EL, Renoir T, Hannan AJ. Environmental enrichment enhances cognitive flexibility in C57BL/6 mice on a touchscreen reversal learning task. Neuropharmacology. 2017 May 1;117:219–226. doi: 10.1016/j.neuropharm.2017.02.009. Epub 2017 Feb 11. PMID: 28196627.

